# Combined strategy of siRNA and osteoclast actin cytoskeleton automated imaging to identify novel regulators of bone resorption shows a non-mitotic function for anillin

**DOI:** 10.1101/435297

**Authors:** Justine Maurin, Anne Morel, Cedric Hassen-Khodja, Virginie Vives, Pierre Jurdic, Irma Machuca-Gayet, Anne Blangy

**Affiliations:** CRBM, Montpellier Univ., CNRS, France; MRI, BioCampus Montpellier, CNRS, INSERM, Univ Montpellier, Montpellier, France; Institut de Génomique Fonctionnelle de Lyon, CNRS UMR3444. Université de Lyon, Ecole Normale Supérieure de Lyon, Lyon, France.; INSERM Unit 1033 and Université Claude Bernard Lyon 1, Lyon, France.

**Keywords:** Osteoclast, Cytoskeleton, Actin, Anillin, Bone resorption, Automated imaging

## Abstract

Osteoclasts are the main cells responsible for the resorption of mineralized extracellular matrices. They are the major targets for anti-resorptive therapies to manage osteoporosis, a major public health problem. Osteoclasts are giant multinucleated cells that can organize their a unique adhesion structure based on a belt of podosomes, which is the keystone of the bone resorption apparatus. We combined differential transcriptomics and siRNA screening approaches to get a broader view of cytoskeletal regulators that participate in the control of osteoclast cytoskeleton and identify novel regulators of bone resorption by osteoclasts. We identified 20 new candidate regulators of osteoclasts cytoskeleton including Fkbp15, Spire1, Tacc2 and RalA, for which we confirmed they are necessary for proper organization of the podosome belt. We also showed that Anillin, well known for its role during cytokinesis, is essential in osteoclasts for correct podosome patterning and efficient bone resorption. In particular, Anillin controls the levels of the GTPase RhoA, a known regulator of osteoclast cytoskeleton and resorption activity. Finally, we set up and validated an automated imaging strategy based on open-source software for automatic and objective measurement of actin cytoskeleton organization in osteoclasts. We provide these pipelines that are useful to automatically assess the effect of collections of siRNAs or chemical compounds on osteoclast cytoskeleton or differentiation.

## Introduction

Osteoclasts have the unique capacity to resorb the mineralized extracellular matrices of the body such as bone and dentin (Cappariello et al., 2014). The highly dynamic process of bone remodeling ensures skeleton maintenance throughout life; it is mediated by a balance between bone resorption by osteoclasts and bone formation by osteoblasts. Unfortunately with age and sex hormone decline as well as in various pathological situations such as bone metastases and inflammatory diseases, osteoclast activity increases and bone formation by osteoblasts becomes insufficient to balance bone degradation. This drives progressive systemic or localized bone loss, leading to osteoporosis, bone frailty and pain that can ultimately cause fractures and impotencies (Seeman, 2003). Targeting the osteoclasts is the main therapeutic strategy to overcome pathological bone loss (Baron et al., 2011). Thus, understanding the molecular mechanisms controlling osteoclast biology is of major importance to develop therapeutic strategies prevent osteoporosis, a major public health problem worldwide (Kastner et al., 2018).

Osteoclasts are multinucleated cells that arise from the fusion of mononucleated hematopoietic precursors. Their differentiation is mediated by two essential cytokines: macrophage colony-stimulating factor (M-CSF) and receptor activator of NFκB (RANKL) (Teitelbaum and Ross, 2003). Osteoclast differentiation provides the cell with the capacity to organize a unique adhesion structure: a belt of podosomes stabilized by microtubules and supporting the ruffled border of the bone resorption apparatus. Within this ring-shape structure, the osteoclast secretes protons, to acidify the extracellular medium and dissolve bone hydroxyapatite, and proteases to digest bone proteins (Touaitahuata et al., 2014). We reported recently that targeting osteoclast cytoskeleton to prevent proper podosome organization is a relevant strategy to prevent pathological bone loss (Vives et al., 2015).

Podosomes are actin-based adhesion units and various regulators of actin dynamics were found to control the bone resorption activity of osteoclasts (Dan Georgess et al., 2014; Touaitahuata et al., 2014) including Rac1 and Rac2 (Croke et al., 2011; Wang et al., 2008) and their exchange factors Vav3 and Dock5 (Faccio et al., 2005; Vives et al., 2011), the GTPases RhoA (Chellaiah et al., 2000) and RhoE/Rnd3 (D. Georgess et al., 2014), the adaptor protein Tensin3 (Touaitahuata et al., 2016) and the actin severing protein Cofilin (Blangy et al., 2012; Zalli et al., 2016). Nevertheless, the entire molecular scheme of actin cytoskeleton regulation required to adequately support bone resorption by osteoclasts is still largely incomplete.

Here, we combined differential transcriptomics and siRNA screening approaches to get a broader view of cytoskeletal regulators that participate in the bone resorption function of osteoclasts and identified novel regulators of bone resorption by osteoclasts. We found that Anillin regulates bone resorption, in particular by impacting on the expression of the GTPase RhoA. We also set up a pipeline to envision larger scale quantitative screening for siRNAs or chemical compounds that affect osteoclast actin cytoskeleton organization and identify new candidate genes involved in bone resorption or candidate anti-resorptive drugs.

## Material and methods

### Mice

Four-week-old C57Bl/6J were purchased from Harlan France and maintained in the animal facilities of the CNRS in Montpellier, France. Procedures involving mice were performed in compliance with local animal welfare laws, guidelines and policies, according to the rules of the regional ethical committee.

### Osteoclast differentiation and siRNA transfection

Primary osteoclasts were obtained from mouse bone marrow as described previously (Touaitahuata et al., 2016; Vives et al., 2011). Briefly, bone marrow macrophages (BMMs) were obtained from long bones of 6- to 8-week-old C57BL/6J mice by growing non-adherent cells for 48h in α-MEM (Lonza) supplemented with 10% heat-inactivated fetal calf serum (FCS, Lonza) and 2 mM glutamine (Lonza) and 30 ng/ml M-CSF (Peprotech). Osteoclasts were then differentiated by culturing BMMs in α-MEM supplemented with 10% heat-inactivated FCS and 2 mM glutamine with 50 ng/ml RANKL (Miltenyi) and 30 ng/ml M-CSF (Peprotech) for 5 days. For siRNA treatments, BMMs were seeded 5×10^5^ cells per well in 24 well plates and transfected for 3 hours at days 2 of osteoclastic differentiation using siImporter (Millipore) with 2.5 pmol siRNA per well in OptiMEM (Thermofisher) complemented 50 ng/ml RANKL and 30 ng/ml M-CSF. The siGenome SmartPool siRNAs were from Dharmacon as described in Supplementary Table 1. Individual siRNAs were purchased as siRNA duplexes from Dharmacon for Spire1, FKBP15, Rala and Tacc2 (Supplementary Table 1) or from Eurogentec: Cy5-labeled negative control siRNA targeting luciferase 5’-CUUACGCUGAGUACUUCGA-3’ according to (Knoch et al., 2004); positive control siRNA targeting mouse Dock5, 5’-GGAGCUCACAAACACGCUACG-3 according to (Touaitahuata et al., 2016); siRNA targeting mouse Anilline 5’- GUAAAGAAAGUCCGUCUU-3’ according to (Lee et al., 2016).

**Table 1:**
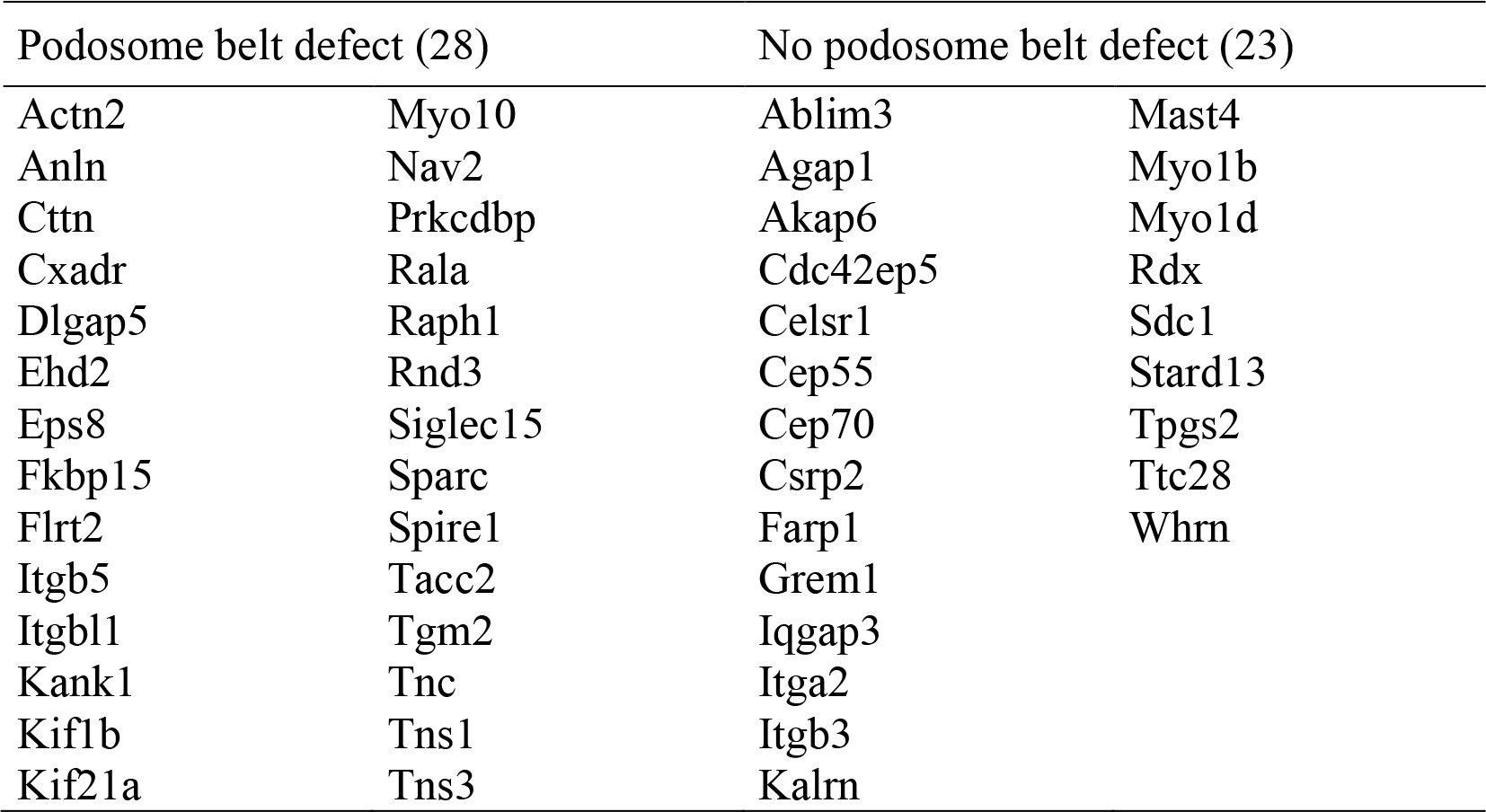
siRNAs pools which, in both screens, provoked podosome belt defects or not.

### Antibodies

Primary antibodies used in this study were as follows, with dilution for western blot (WB) and immunofluorescence (IF): anti-α-Tubulin T5168 (Sigma) 1:5000 (WB), 1:2000 (IF with Arrayscan VTi microscope), anti-α-Tubulin (YOL 1/34) sc-53030 (Santa Cruz Biotechnology), 1:100 (IF), anti-vinculin V9131 (Sigma) 1:4000 (WB), 1:400 (IF), anti-Phospho-Vinculin (Y1065) 44-10786 (Invitrogen) 1:1000 (WB), anti-Phospho-MLC (Thr18/ser19) ref 3674 (Cell signaling) 1:1000 (WB), 1:50 (IF), anti-MLC (clone MY21) M4401 (Sigma) 1:1000 (WB), anti-cofilin sc-376476 (Santa Cruz Biotechnology) 1:1000 (WB), anti-Phospho-cofilin (ser3) sc-12912 (Santa cruz) 1:500 (WB), anti-acetylated α- Tubulin (clone 611B1) T6793 (Sigma) 1:1000 (WB), anti-RhoA (clone 26C4) sc-418 (Santa Cruz Biotechnology) 1:500 (WB), 1:50 (IF), anti-MyosinIIA PRB-440P (Covance) 1:500 (IF), anti-Anillin GTX 107742 (Genetex) 1:100 (IF). For immunofluorescence, Alexa-labeled antibodies (Life Technologies) were all used 1:1000: donkey Alexa488-anti-mouse antibody A21202, donkey Alexa546-anti-mouse antibody A10036, donkey Alexa488-anti-rabbit antibody A21206, goat Alexa633-anti-rat antibody A21094.

### Immunofluorescence

At the end of differentiation, osteoclasts were fixed for 20 minutes in 10 μM Taxol (Sigma) and 3.2% paraformaldehyde in PHEM (60 mM Pipes, 25 mM Hepes, 10 mM EGTA, 4 mM MgSO4, pH 6,9). After permeabilization with 0.1% Triton X-100 in PBS for 1 minute and blocking with 1% BSA in PBS, osteoclasts were stained for one hour with relevant primary antibodies followed by one hour with fluorescent secondary antibody and if appropriate Alexa564-Phalloidin A22283 or Alexa647-Phalloidin A22287 (Life Technologies) 1:1000, to stain filamentous actin. When necessary, DNA was stained 15 minutes with bisbenzimide Hoechst dye (Sigma). For manual confocal and wide field microscopy, coverslips were mounted in CitiFluor mounting medium (Biovalley) images were acquired with manually with a Leica DM6000 wide field microscope using a 63X HCX Plan 0.6-1.32 NA oil (designed thereafter as high magnification) objective or with a Leica SP5 confocal microscope using a 63X HCX Plan Apo CS oil 1.4NA objective. For automated imaging (24-well plates), images were acquired with an automated Arrayscan VTi microscope (Thermo) equipped with a 10X EC Plan Neofluar 0.3NA (designed thereafter as low magnification) objective. Each well was imaged as a spiral mosaic of about 300 partially overlapping fields at 3 wave lengths: 349 nm for Hoechst, 488 nm green for tubulin and 540 nm for actin. Illumination was corrected in each image with CellProfiler software 3.0 (see below) and then, they were combined in ImageJ 1.51w to obtain an image of the whole well. All images where then processed using ImageJ 1.51w or Adobe Photoshop CS5 or CS6 and mounted with Adobe Illustrator CS5 or CS6. All imaging was performed at the Montpellier RIO Imaging facility (http://www.mri.cnrs.fr).

### Automated segmentation of osteoclast and peripheral actin intensity measurement

Images were analyzed using custom pipelines we built in the freely available CellProfiler 3.0 software (Carpenter et al., 2006), as described in Supplementary Tables 2 and 3. Briefly, we corrected each original Arrayscan VTi images in the 3 illumination channels for background illumination heterogeneities introduced by the microscope optics with CorrectIlluminationCalculate module, before combining the images in ImageJ 1.51w to obtain one image per well. To perform objects identification and features extraction, the original images were rescaled to have pixel values between 0 and 1 thanks to the module RescaleIntensity, thereby all images were in the same range and suitable for comparison. The IdentifyPrimaryObjects module was used to find the cells in the tubulin channel and the nuclei in the Hoechst channel. The number of nuclei per cell was obtained with the RelateObjects module. We applied filters with the FilterObjects module for the cell size and number of nuclei per cells. The modules MeasureObjectSizeShape and MeasureObjectIntensity were used to extract osteoclast features. ExpandOrShrinksObjects module was used to define an 8-pixel wide band (with 1 pixel = 1.024 μm) inside the periphery of segmented objects. For object classification and osteoclast selection among the segmented objects, we performed supervised classification using RandmForest under R (Breiman, 2001).

**Table 2:**
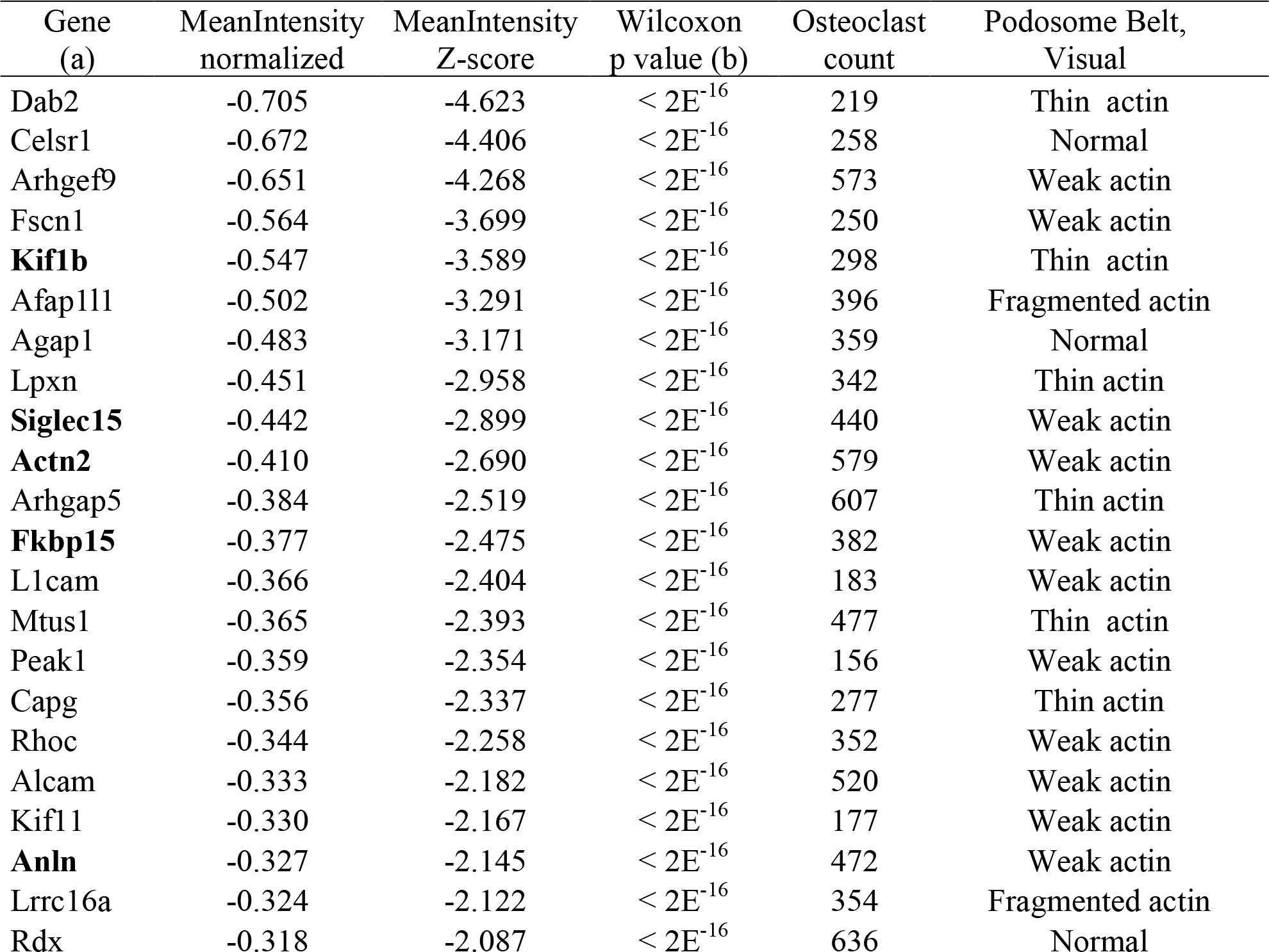
Comparison between automated and visual assessment of actin organization.

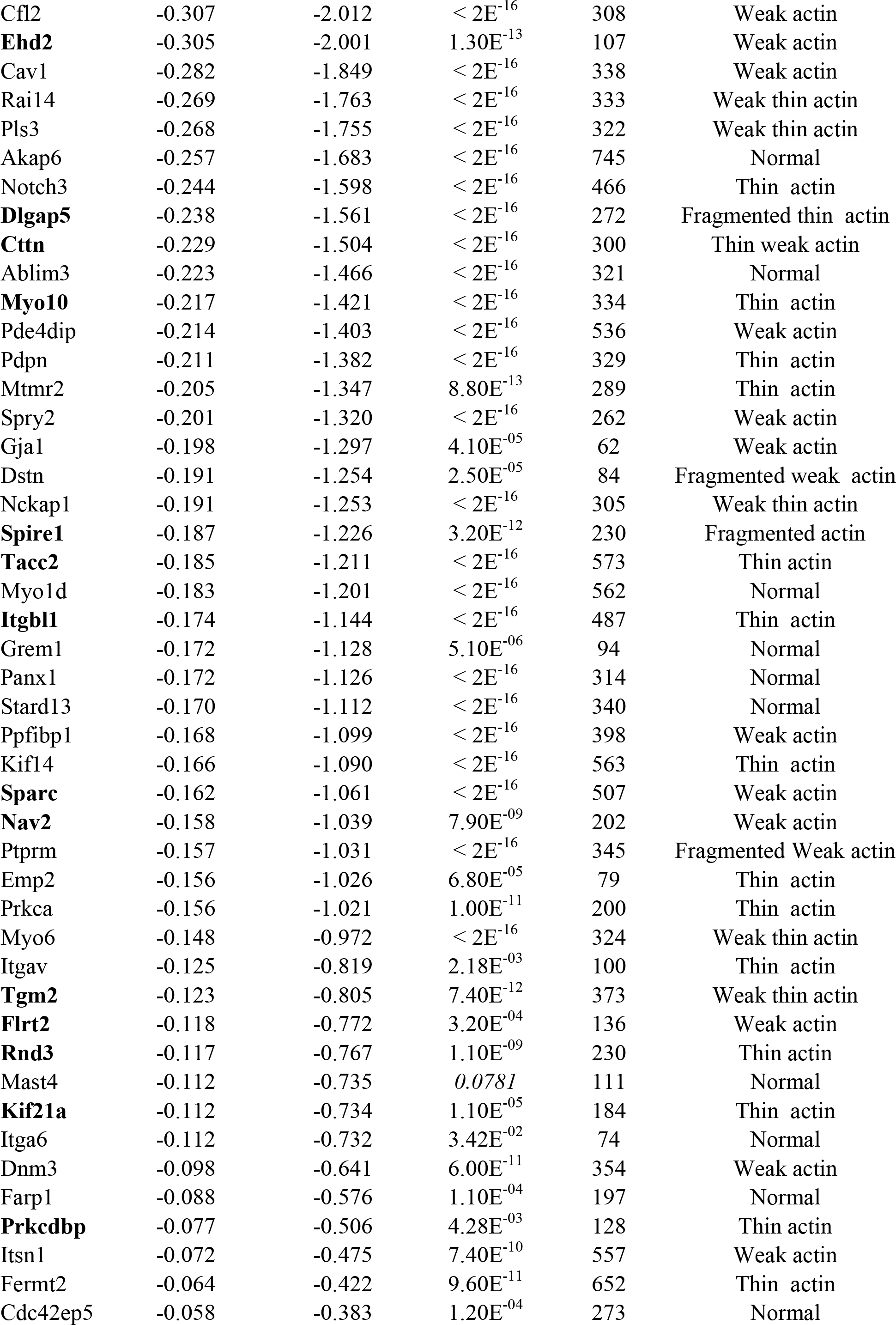

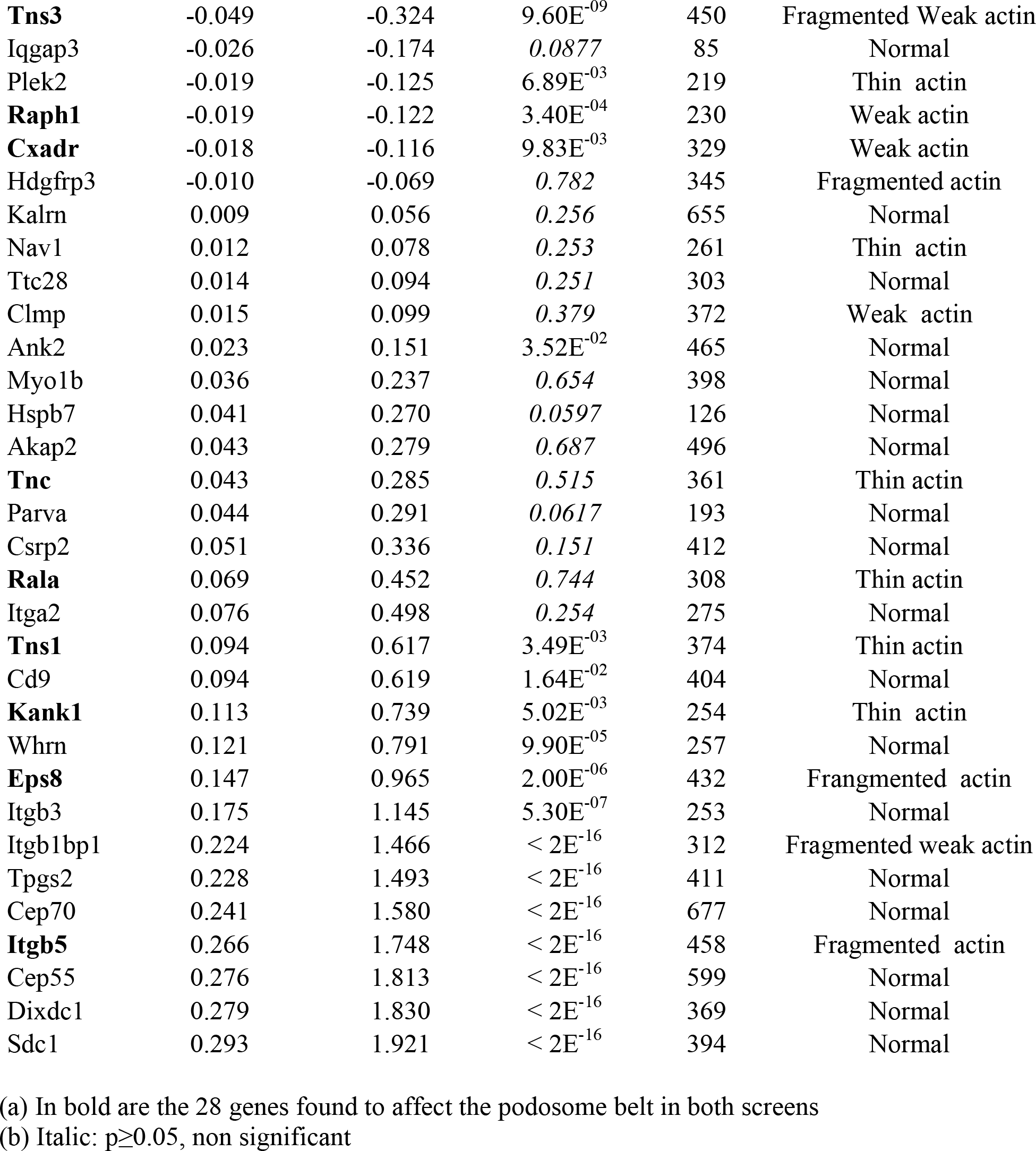

### Western blots

Whole cell extracts were prepared in Laemmli sample buffer, resolved on SDS-PAGE and electrotransferred on nitrocellulose or PVDF membranes. Immunoblotting was performed and signals were revealed either using the ECL Western Lightning Plus detection system (Perkin Elmer) with horseradish peroxidase-conjugated secondary antibodies 1:10,000 anti-mouse NA931V or anti-rabbit NA934V (GE Healthcare).

### Real time Q-PCR

DNaseI-treated total RNA was extracted using the High Pure RNA Isolation kit (Roche) and to generate cDNAs, RNA was primed with 10-mer random primers and reverse transcription was catalyzed using Superscript II reverse transcriptase (Invitrogen). Q-PCR was performed an Mx3000p PCR system (Stratagene) using the Platinium Taq DNA polymerase (Invitrogen) and SYBR Green I (Bio Wittaker) as described (Coelho et al., 2005). Primers used were 5’-ACAGTCCATGCCATCACTGCC-3’ and 5’-GCCTGCTTCACCACCTTCTT-3’ for Gapdh and 5’-TTCGGAATGACGAGCACACG-3’ and 5’-GTCTAGCTTGCAGAGCAGCT-3’ for RhoA (Brazier et al., 2006); 5’-CAGTAGCGAGATCTGCCCCG-3’ and 5’-TCGGAGAACAGGGCTTTGCA-3’ for Anln (Lee et al., 2016). The threshold cycle (Ct) of each amplification curve was calculated by Roche Diagnostics LightCycler 480 software using the second derivative maximum method. The relative amount of a given mRNA was calculated using the ∆Ct method (Livak and Schmittgen, 2001).

### Apatite collagen complexe (ACC)-coated substrate preparation and osteoclast seeding

The ACCs (apatite collagen complexes) were prepared using the method described previously (Saltel et al., 2004). Briefly, 11-mm diameter glass coverslips were coated with calf skin type I collagen (Sigma) and incubated for 7 days at 37°C in 200 mM Tris-buffered saline (TBS) pH 8.5, containing egg yolk phosvitin (0.13 mg/ml), alkaline phosphatase (0.13 mg/ml) and dimethyl suberimidate hydrochloride (1 mg/ml) (Sigma) as a cross-linking reagent. Slides were washed several times with TBS and incubated for 3 hours at 37°C in the same medium without cross-linking reagent. Then, they were incubated for 20 hours at 37°C in 200 mM TBS pH 8.5 containing 6 mM calcium β-glycerophosphate (Sigma). The last 2 steps were repeated 7 to 14 times, depending on the amount of precipitated calcium phosphate. After washing with PBS, the glass slides were dried. Osteoclasts at day 3 of differentiation were detached with Accutase (Sigma) for 5 to 10 minutes at 37°C, seeded and grown for 1 to 2 more days onto ACC before proceeding for immunofluorescence.

### Osteoclast activity assays

Mineral dissolution activity of osteoclasts was measured as described (Touaitahuata et al., 2016). Briefly, at day 3 of differentiation (24 h after siRNA transfection), osteoclasts were rinsed once in PBS, detached with Accutase (Sigma) for 5 to 10 minutes at 37°C and seeded for 3 days onto inorganic crystalline calcium phosphate (CaP)-coated multiwells (Osteo Assay Surface, Corning). In each experiment, four wells were then stained for Tartrate Resistant Acid Phosphatase (TRAP) activity to count osteoclasts and four wells with Von Kossa stain to measure CaP dissolution as described previously (Vives et al., 2011). Von Kossa and TRAP staining were imaged with a Nikon SMZ1000 stereomicroscope equipped with a Nikon DXM 1200F CCD camera. Quantification osteoclasts and resorbed areas were done with ImageJ 1.51w software. Osteoclast specific activity was expressed as the average area resorbed in the 4 wells stained with von Kossa normalized by the average number of osteoclasts in the 4 wells stained with TRAP.

### Statistics

Statistical significance was assessed with the non-parametric statistical tests mentioned in the figure legend. All analyses were done with GraphPad Prism (GraphPad Software, Inc., La Jolla, CA), with p<0.05 considered statistically significant. Unbiased segmentation image analysis was performed with RandomForest method in R open-source software.

## Results

### Selection of potential new regulators of osteoclast cytoskeleton

In view of identifying novel regulators of osteoclast biology, in particular genes involved in the regulation of bone resorption via the control of the cytoskeleton, we adopted a candidate strategy. We took advantage of our former Affymetrix transcriptomic analysis comparing osteoclasts derived either directly from human blood monocytes (Mo-OC) or trans-differentiated from dendritic cells (DC) derived from the same monocytes (DC-OC) (Gallois et al., 2010). In fact, monocytes (Mo), OC and DC are myeloid cells sharing the capacity to assemble podosomes, but only OC (Mo-OC and DC-OC) are able to organize them into a belt to form the bone resorption apparatus. We reasoned that genes induced during both osteoclast differentiation pathways could represent novel regulators of osteoclast biology. We compared the transcriptomic data of Mo, DC, Mo-OC and DC-OC from the 2 donors of this Affymetrix study and selected the U133 Plus 2.0 probesets showing at least 3-fold increase both between Mo and Mo-OC and between DC and DC-OC. Among the 2337 Affymetrix probesets that met these criteria, we eliminated the probesets qualified “Absent” in the two samples of Mo-OC or of DC-OC. This led to 1118 probesets corresponding to 808 unique genes. Then, we checked among the GO terms associated with these genes to identify those involved in the regulation of actin and microtubule cytoskeleton. This led to a second list of 100 genes (Supplementary Table 1), including known regulators of osteoclast cytoskeleton such as Tns3 (Touaitahuata et al., 2016) and RhoE (D. Georgess et al., 2014).

### Selection of candidate regulators of osteoclast podosome belt formation

We set up an siRNA screening approach based on automated imaging to find among the 100-cytoskeleton regulating genes those affecting the organization of podosomes in osteoclasts sitting on plastic. For each gene, we used a commercial SmartPool of 4 siRNAs (Supplementary Table 1), which was transfected at day 2 of osteoclast differentiation from mouse bone marrow macrophages in 24-well plastic plates. Transfection was performed after 2 days of RANKL treatment to minimize the risk of interfering with the early osteoclast-differentiation processes and maximize the chances for the interference to be efficient during osteoclast podosome rearrangements, which occurs mainly during days 3 to 4. Osteoclasts were then grown for 2 more days before fixation, staining for DNA, tubulin and actin. The experiment was done in duplicate with 2 independent mouse osteoclast samples. Entire wells were imaged as a mosaic on an automated microscope with a 10x low magnification objective and full-well images were reconstituted for DNA, actin and tubulin.

We then examined manually the structure of actin, seeking siRNAs affecting the dense peripheral actin ring (Figure 1A), such as the positive control Dock5 siRNA, known to perturb the formation of the podosome belt (Vives et al., 2011). As a negative control, we used Luciferase siRNA, which does not target any mouse gene and should not affect the podosome belt (Vives et al., 2011). In negative control wells transfected with Luciferase siRNA, low magnification imaging of actin clearly reveals the dense podosome belt at the periphery of the osteoclasts; in contrast, actin staining appears diffuse in the positive control wells transfected with Dock5 siRNA (Figure 1B). We defined the siRNAs affecting the podosome belt as siRNAs that caused actin staining to become fragmented and/or weaker and/or thinner and/or absent, as exemplified in the bottom panels of Figure 1A, in more than half of the osteoclast periphery and in the majority of osteoclasts, provided that this was a visual estimation and not a quantification.

**Figure 1:**
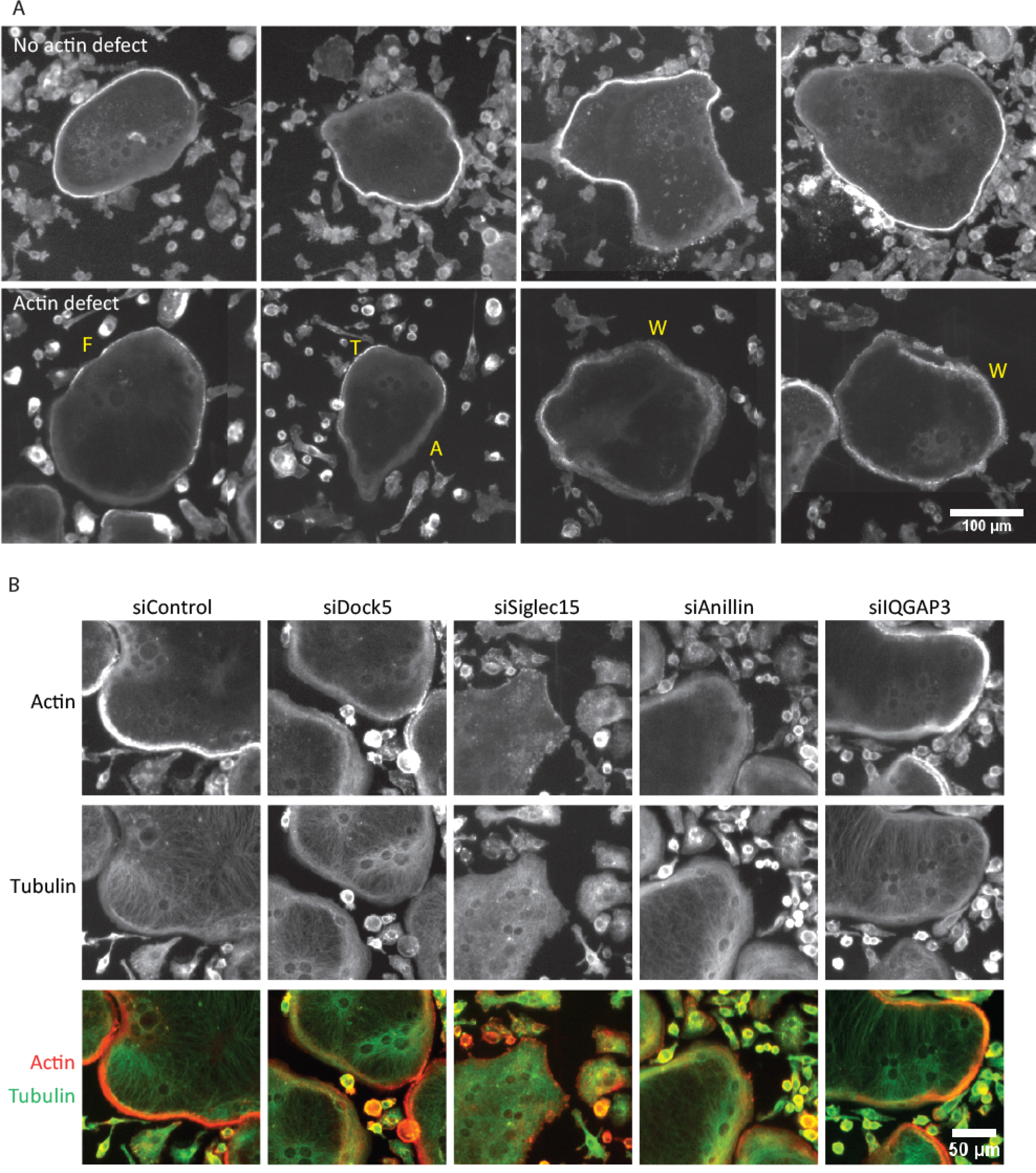
Selection of candidate regulators of osteoclast actin cytoskeleton. Automated Arrayscan VTi microscope images of osteoclasts plated in plastic 24-well plates showing (A) actin staining in representative osteoclasts after siRNA transfection to exemplify different podosome belt phenotypes: normal podosome belts (to panels) and abnormal podosome belts (bottom panels): F= Fragmented, T=Thin, W=Weak, A=Absent; and (B) actin staining (top panels) and tubulin staining (middle panels) overlaid in the bottom panels, in representative fields after transfection of Luciferase negative control (siControl), Dock5 positive control (siDock5) siRNAs and test siRNA SmartPools (siSiglec15, siAnillin and siIQGAP3).

Among the 100 SmartPool siRNAs tested, 28 were found to perturb the peripheral actin staining in both experiments, suggesting that the target genes could participate in the formation of the podosome belt (Table 1). These included the known regulators of osteoclast actin cytoskeleton Siglec15 (Ishida-Kitagawa et al., 2012) that also appears to perturb the organization of microtubules (Figure 1B), Tns3 (Touaitahuata et al., 2016) and RhoE (D. Georgess et al., 2014) (Table 1). Conversely, 23 siRNA pools, such as IQGAP3 (Figure 1B), did not provoke any disturbance of actin staining in both screen (Table 1), suggesting either that gene silencing was not efficient or that the target genes, although related to the control of actin dynamics, do not affect the podosome belt when silencing is performed at day 2 of differentiation.

Among the 100 candidate genes selected upon differential transcriptomics, siRNA screening highlighted 28 genes as potential regulators of actin organization in osteoclasts, of which 22 are of unknown function in osteoclasts as yet.

### Fkbp15, Spire1, RalA and Tacc2 are necessary for correct podosome belt formation

To validate our result, we first chose 4 genes of the 22 of unknown function in osteoclasts that affect actin organization in our screen: Fkbp15, Spire1, Tacc2 and RalA. We previously found Fkbp15 as a major partner of Dock5 (Touaitahuata et al., 2016), an exchange factor for Rac essential for osteoclast actin organization and bone resorption (Vives et al., 2011). Spire1 is a regulator of actin associated with invadopodia and involved in cancer cell invasion (Lagal et al., 2014). Tacc2 is a microtubule +TIP protein promoting microtubule growth that localizes in front of EB1 (Rutherford et al., 2016), another +TIP essential for the formation of osteoclast podosome belt (Biosse Duplan et al., 2014). Finally, the small GTPase RalA is an important regulator of exocytosis (van Dam and Robinson, 2006), a basal process for osteoclast function. For each gene, the 4 siRNAs from the SmartPool used in the screen were individually transfected in osteoclasts differentiated on glass coverslips and organization of the podosome belt was analyzed by imaging with a high magnification 63x objective after actin and vinculin fluorescent staining (Figure 2A). For the 4 genes, siRNAs provoked an important reduction of the proportion of cells with a properly organized podosome belt (Figure 2B). We observed that the podosomes were not condensed in more than half of the osteoclast periphery: individual podosome cores were visible by actin staining and/or a ring of vinculin was surrounding the podosome core, contrarily to the Luciferase siRNA control, in which the belt is made of highly condensed podosomes where actin cores are not individualized (Figure 2A). Of note, tubulin staining did not reveal any effect of Tacc2 siRNAs on the organization of microtubules in fixed osteoclasts (data not shown).

**Figure 2:**
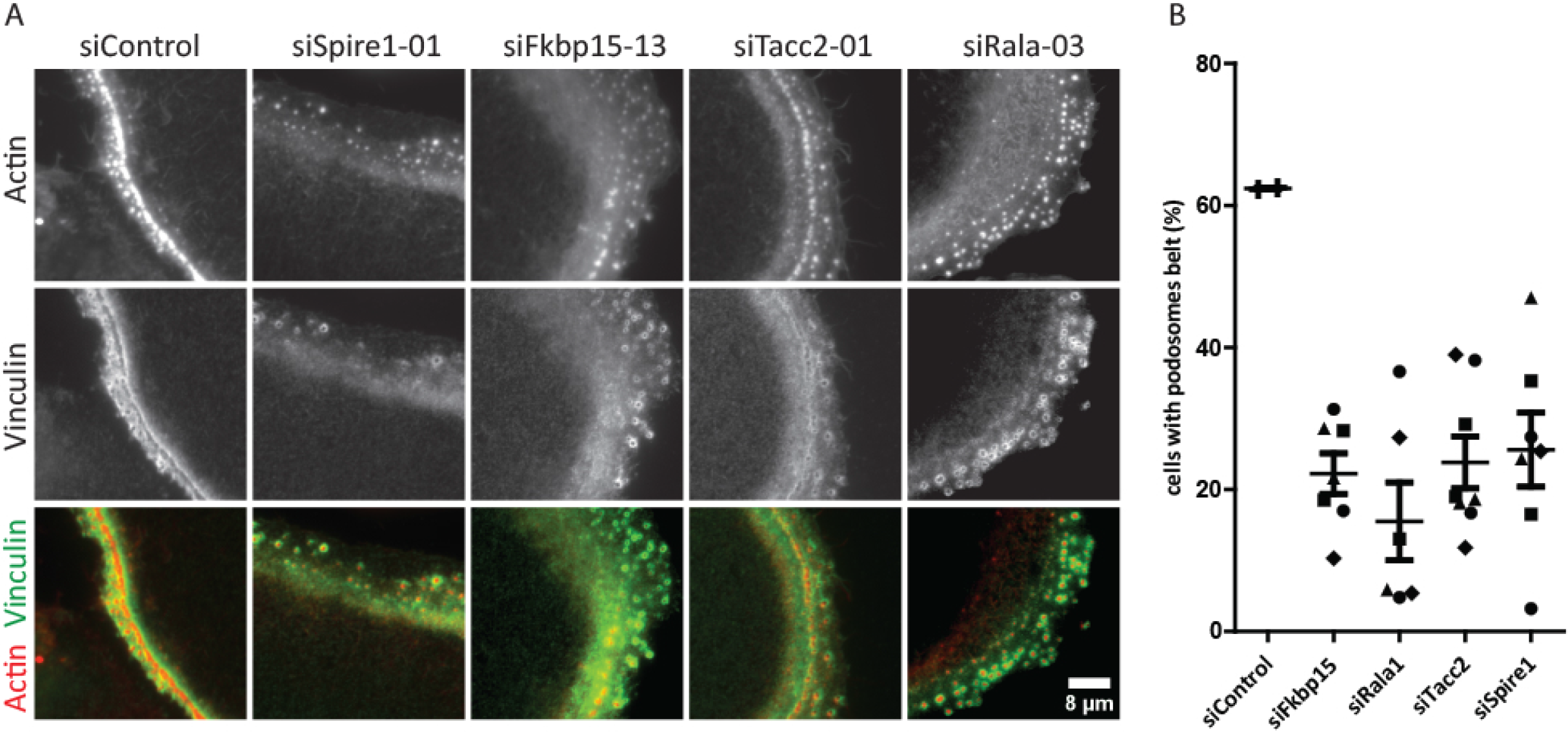
Importance of Fkbp15, Spire1, RalA and Tacc2 for correct of the podosome belt organization. (A) Wide field image showing a representative podosome belt in osteoclast sitting on glass and transfected with after Luciferase negative control (siControl) siRNA or with one siRNA directed against Fkbp15, Spire1, RalA and Tacc2; the siRNA numbers refer to the duplex catalog numbers in Supplementary Table 1. Osteoclasts were labeled for vinculin and actin. (B) Bar graph Graph showing average and SEM percent of osteoclasts with a podosome belt after transfection of Luciferase (siControl) siRNA or the 4 different siRNAs targeting Fkbp15 (square: siRNA 13, triangle: 14, diamond: 15, dot: 16 according to Supplementary Table 1), Spire1, RalA and Tacc2 (square: siRNA 13, triangle: 14, diamond: 15, dot: 16 according to Supplementary Table 1), from two independent experiments counting over 100 osteoclasts per experiments and per siRNA.

These results show that among the 22 genes of unknown function in osteoclasts that we highlighted in our siRNA screen, there are indeed novel regulators of osteoclast cytoskeleton.

### Anillin regulates osteoclast podosome belt and bone resorption

Our screen revealed severe defect in actin organization in osteoclasts receiving Anillin SmartPool siRNA (Figure 1B). Anillin is essentially known as a key protein during cytokinesis (Zhang and Maddox, 2010), which coordinates RhoA and Myosin II activities at the contractile ring midzone (Piekny and Glotzer, 2008; Straight et al., 2005) and binds to microtubules to properly organize the mitotic spindle (van Oostende Triplet et al., 2014). In osteoclasts plated on glass coverslips, we found that Anillin localizes at the podosome belt (Figure 3A and Supplementary Figure 1A). Similar to other components of the podosome belt such as vinculin, Anillin staining at osteoclast periphery appeared more diffuse where the podosome belt was not present, as illustrated at the top left of the cell shown in Supplementary Figure 1A. This is consistent with Anillin being associated with the podosome belt. Consistent with previous studies demonstrating nuclear localization of Anillin, as referenced in (Wang et al., 2015), we also found Anillin accumulated in the nucleus of osteoclasts (Supplementary Figure S1A). We further analyzed the localization of Anillin in osteoclasts sitting on the mineralized apatite collagen complex-coated substrate ACC (Figure 3B-D and Supplementary Figure 1B-D). In early forming sealing zones, where actin organizes as a small circular shape, Anillin localizes as a thin ring in the inner part of the actin ring (Figure 3B and Supplementary Figure 1B) whereas Vinculin rather colocalizes with actin. In more mature circular sealing zones (Figure 3C and Supplementary Figure 1C), the distribution of Anillin gets broader inside the actin ring whereas Vinculin remains at the ring. Finally, disassembling sealing zones that exhibit more irregular shape, Anillin spreads inside the center of the actin ring (Figure 3D and Supplementary Figure 1D).

**Figure 3:**
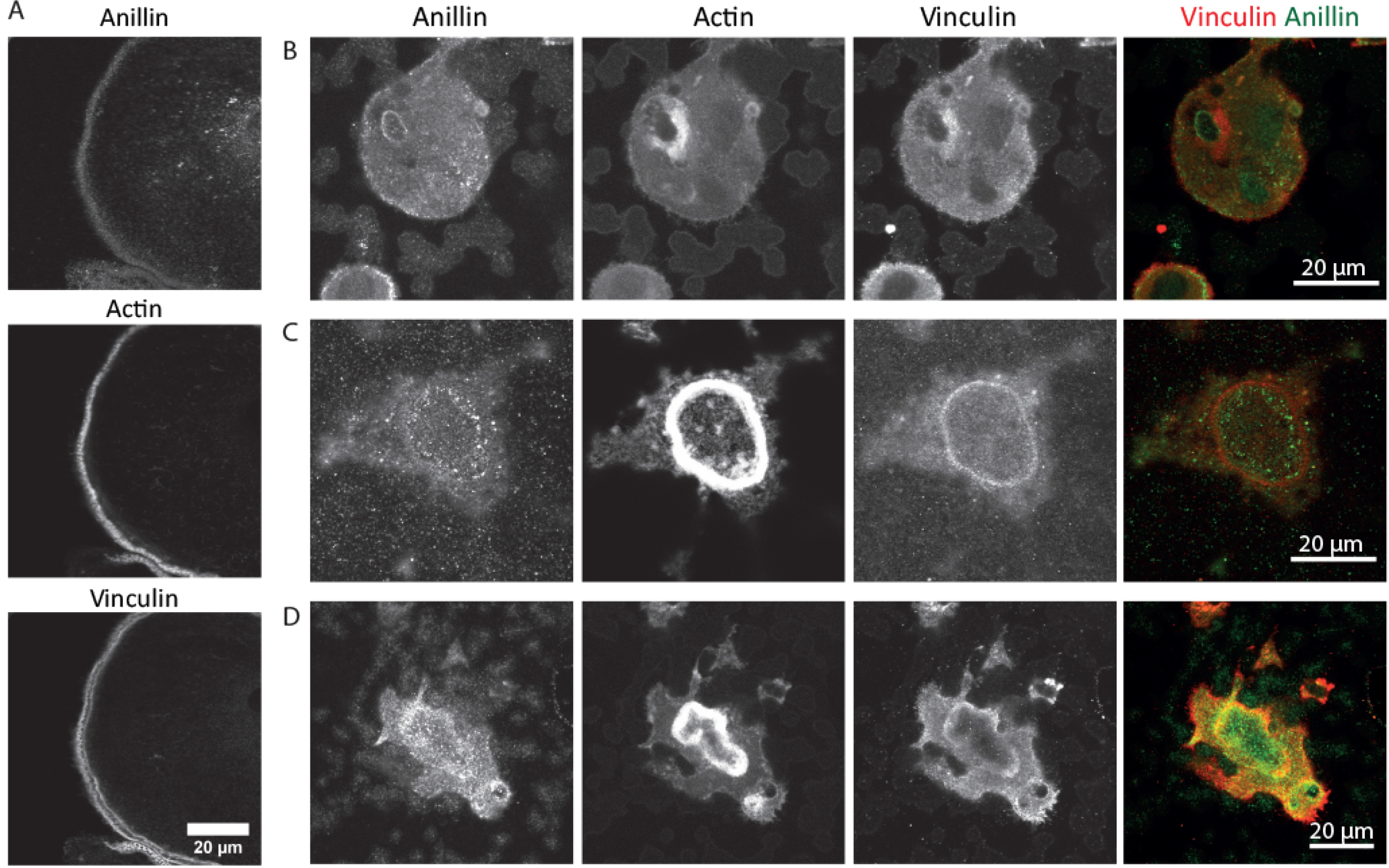
Loclaization of Anillin at the podosome belt and at the sealing zone. (A) Wide field image showing Anillin localization at the podosome belt in a representative osteoclast sitting on glass and labeled with vinculin and actin. Confocal images are also shown in Supplementary Figure 1A. (B-D) Maximum z projection of 3 consecutive confocal plans, with 170 nm increment between planes, showing Anillin localization at the sealing zone of osteoclasts sitting on ACC substrate and labeled with actin and vinculin, and showing an early forming sealing zone (B), a mature sealing zone (C) and a late sealing zone (D). The individual plans in image gallery are shown in Supplementary Figure 1B-D

As thigh control of RhoA levels is essential for proper organization of osteoclast podosomes and for bone resorption (Ory et al., 2008) and Myosin IIA localizes at the podosome belt (McMichael et al., 2009), we explored a potential non-mitotic role of Anillin in osteoclasts. We first confirmed the siRNA screen results by silencing Anillin with an individual siRNA formerly validated in mouse oocytes (Lee et al., 2016). This Anillin siRNA efficiently diminished Anillin mRNA levels (Figure 4A) and provoked actin disorganization in osteoclasts plated on glass: podosomes were located at the cell periphery but they were improperly condensed (Figure 4B-C), thus confirming the results obtained in the siRNA screen with the SmartPool. Similarly on ACC, we found that the fraction of osteoclasts with a podosome belt was strongly reduced upon treatment with Anillin siRNA, as compared to control Luciferase siRNA (Figure 4D and Supplementary Figure 2A). We then examined the consequence of Anillin silencing on osteoclasts activity. Consistent with the severe disorganization of the podosome belt, transfection of Anillin siRNA strongly reduced the mineral dissolution activity of osteoclasts as compared to control Luciferase siRNA (Figure 4E and Supplementary Figure 2B).

**Figure 4:**
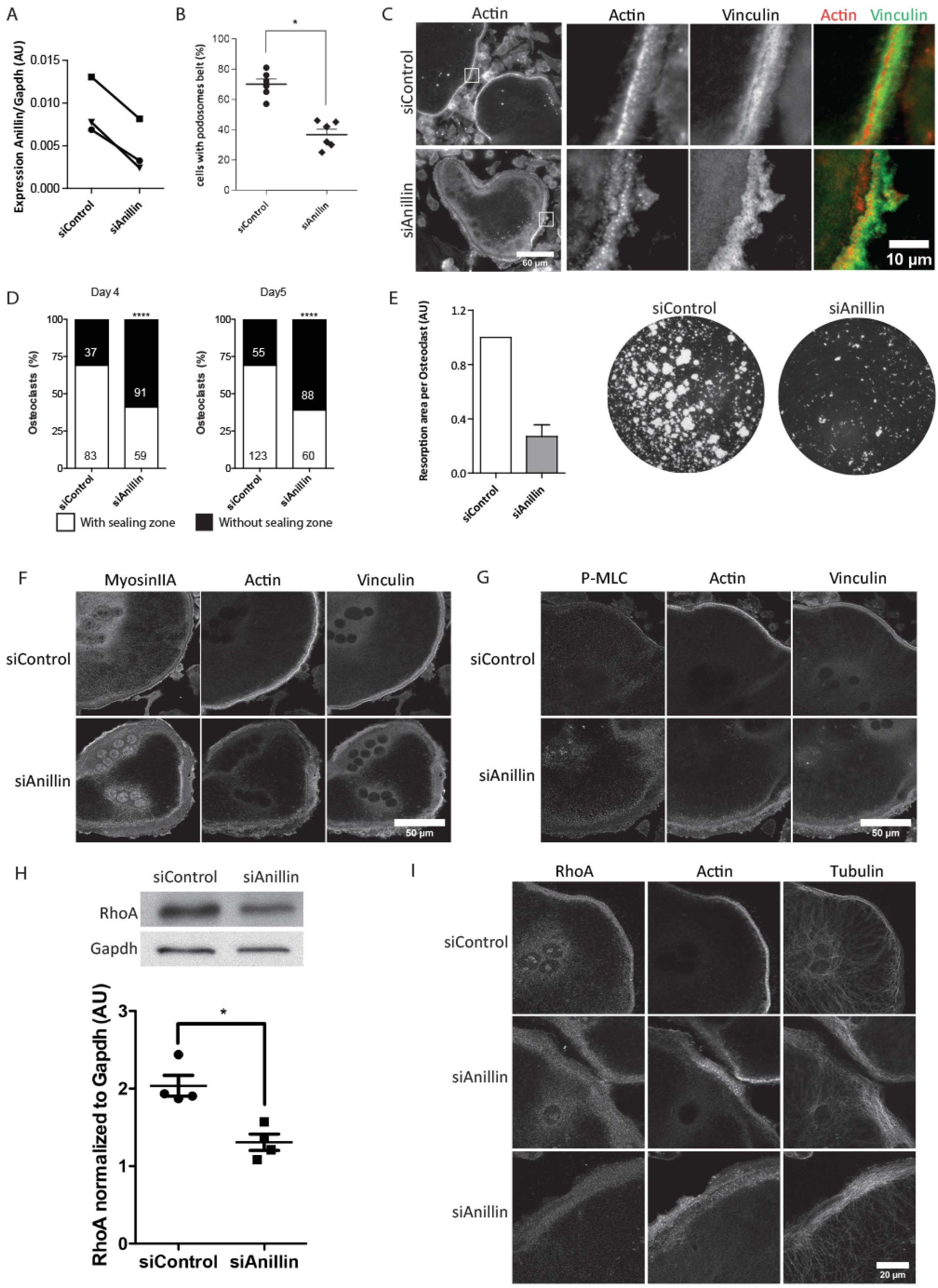
Silencing of Anillin disorganizes the podosomes, inhibits osteoclast activity and decreases the total level of RhoA in osteoclasts. (A) Bar graph Graph comparing AnillinmRNA levels normalized to Gapdh in osteoclasts transfected with control luciferase siRNA (siControl) or Anillin siRNA (siAnillin), in three independent experiments and measured by QPCR. (B) Bar graph Graph showing average and SEM percent of osteoclasts with a normal podosome belt after siControl or siAnillin transfection, from six independent experiments counting over 100 osteoclasts per experiments, *p<0.05 (Mann’Whitney test). (C) Wide field image showing a representative normal podosome belt after siControl treatment or a disorganized podosome belt after siAnillin treatment, in osteoclasts sitting in glass and labeled for vinculin and actin. (D) Graphs comparing the proportion of osteoclasts seeded on ACC and presenting a sealing zone at Day 4 and Day 5 of differentiation, after siControl or siAnillin transfection at Day 2. Figures indicate the number of osteoclasts counted in each category. White: with sealing zone, Black: without sealing zone; **** p<0.0001 Fisher’s exact test. (E) Bar graph showing resorption area per osteoclast treated with siControl or siAnillin, determined after von Kossa staining of calcium phosphate matrices (representative well images below) and normalized to osteoclast numbers determined after TRAP staining, with siControl normalized to 1 in n=4 independent experiments. Individual experiment results are shown in Supplementary Figure 2. (F-G-I) Single z-plan confocal images of representative osteoclasts sitting on glass, showing (F) MyosinIIA, (G) P-MLC and (I) RhoA localization in siControl and siAnillin treated-osteoclasts and labeled for actin and vinculin. (H) Representative western blot and quantification of the level of RhoA protein in osteoclasts transfected with siControl or siAnillin. Graph shows the levels of RhoA protein normalized to the levels of Gapdh, from four independent experiments; * p<0.05 % Mann Whitney test.

We further examined which signaling pathways could be affected by Anillin depletion in osteoclasts. In osteoclasts plated on glass, MyosinIIA and phosphorylated myosin light chain (P-MLC) remained at the cell periphery (Figure 4F-G) and western blot showed that the global level of MLC and P-MLC were not affected by Anillin siRNAs (Supplementary Figure 3). Similarly, we did not find by western blot any significant change in the levels of total and acetylated tubulin, total and phosphorylated vinculin and cofilin (Supplementary Figure 3), two proteins important for podosome belt organization and for bone resorption (Blangy et al., 2012; Fukunaga et al., 2014; Zalli et al., 2016). Interestingly, we repeatedly found that Anillin siRNAs led to a reduction of total RhoA (Figure 4H), while protein accumulation at the periphery of osteoclasts sitting on glass was not affected (Figure 4I). Conversely, the levels of RhoA mRNA were not affected by Anillin depletion (data not shown). Of note, we did not find any obvious effect of Anillin on microtubule organization in fixed osteoclasts, neither with the smartpool siRNAs (Figure 1B) nor with the individual siRNA (Figure 4I).

This suggests Anillin is important for osteoclast function by it role in actin cytoskeleton organization, which involves the control of GTPase RhoA levels and unknown downstream effectors.

### Automated detection of osteoclasts and peripheral actin quantification

In our study, we applied a targeted approach to identify novel regulators of bone resorption. Although successful, this approach is biased by the fact that the very time consuming observations of osteoclasts must be done by eye and that thus initial selection criterions are qualitative. In order to envision less-biased and higher-content screening for siRNAs or molecules interfering with osteoclast cytoskeleton, we sought to set up an automated method to quantify actin organization in osteoclasts. We based our development on the images and data from one of our 100-gene siRNA screens.

In a first intension, we developed a pipeline to automatically segment osteoclasts in the complete well images with CellProfiler pipelines (Supplementary Tables 2 and 3). CellProfiler IlluminationCorrection module was applied to compensate for the non-uniformities in field illumination in each image (Figure 5A) before stitching to create of whole well images (Supplementary Table 2). The original image fluorescence was rescaled to have pixel values between 0 and 1 in the 3 channels thanks to the module RescaleIntensity (Supplementary Table 3). We performed automated segmentation of cells in the tubulin and of nuclei in the Hoechst channel respectively, using the IdentifyPrimaryObjects module of CellProfiler (Figure 5B-C). We obtained the number of nuclei per cell with the RelateObjects module. To select osteoclasts and remove the mononucleated cells, we applied 2 filters with the FilterObjects module: the first filter relates to cell size, and excludes cells with surface less than 4000 pixels and the second relates to the number of nuclei per cells, and only retains cells that contain at least 3 nuclei (Supplementary Table 3 and orange contour cells in Figure 5D). Finally, using IdentifyTernaryObjects in CellProfiler (Supplementary Table 3), we defined an 8-pixel wide peripheral ring inside each object, which contains the podosome belt if present (Figure 5D). In the actin channel, we then measured the mean intensity of actin staining within this ring, a parameter expected to reflect the organization of the podosome belt.

**Figure 5:**
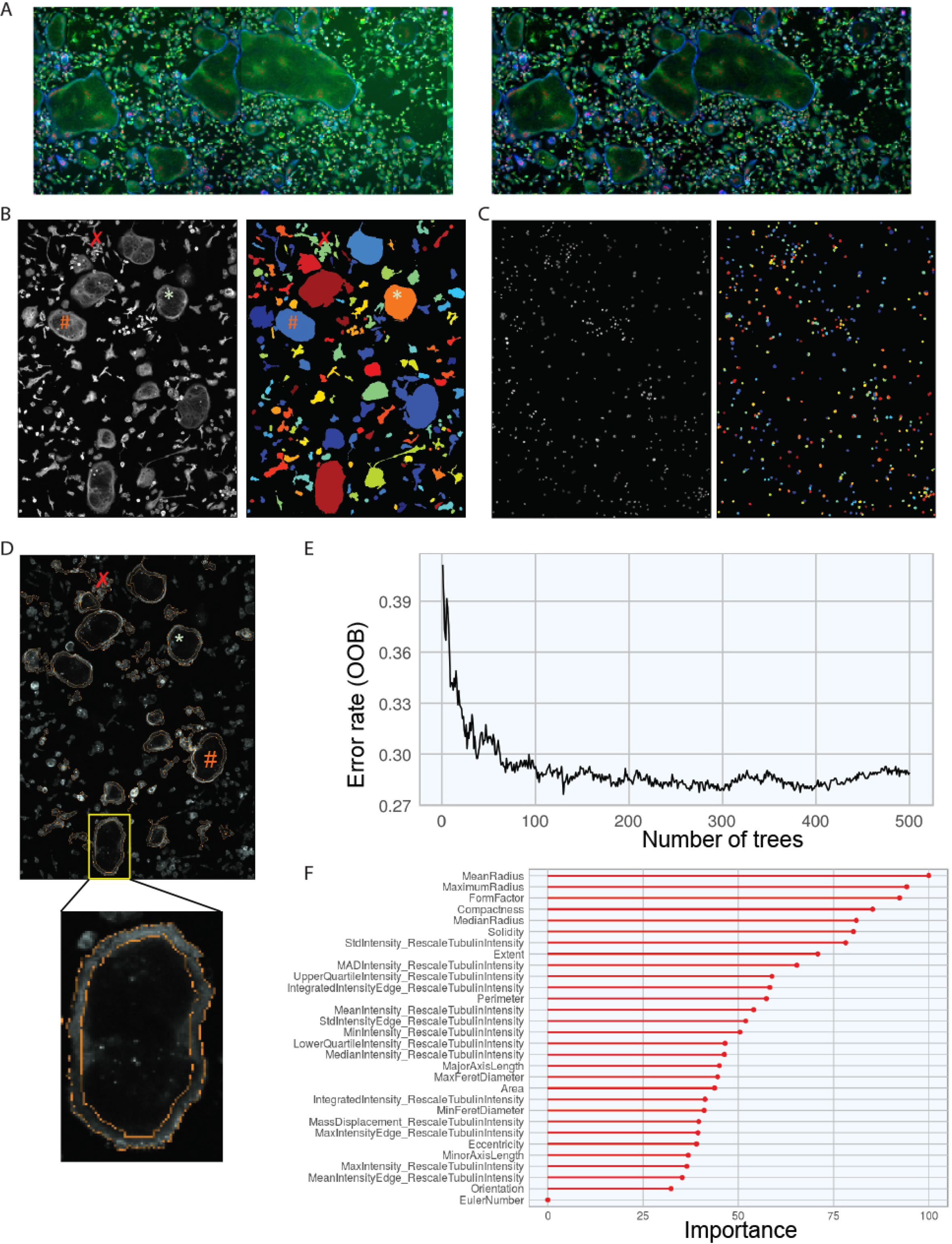
Automated selection of osteoclasts using CellProfiler and RandomForest. (A) IlluminationCorrection module: the images show a representative area of the global image of a control Luciferase siRNA transfected well, obtained after assembling individual Arrayscan VTi microscope images taken with a 10x magnification objective, uncorrected (left) or corrected (right) for non-homogeneous illumination of the individual fields. (B-C) IdentifyPrimeryObject module: the images show (B) cell segmentation in the tubulin channel and (C) nuclei segmentation in the Hoechst channel, in the same representative area of the global image of a control Luciferase siRNA transfected well. (D) RelateObjects, FilterObjects and IdentifyTertiaryObject modules: in the image shown in (B-C), selection of objects with an area greater than 4000 pixels and containing at least 3 nuclei (orange line surrounded cells) and definition of an 8-pixel wide peripheral ring inside each object, expected to contain the podosome belt in the actin channel (magnified area), with 1 pixel = 1.024 μm. In (B-D), *, # and X symbols respectively show examples of properly segmented osteoclasts, osteoclasts surrounded by a few mononucleated cells and mononucleated cell aggregates not containing osteoclasts. (E) Plot showing the measurement of the object classification error rate at the addition of each new tree during the training phase of RandomForest and the convergence to a 0.28 value of the error rate. (F) Respective parameter importance for the prediction of object classes with RandomForest.

To assess the accuracy of osteoclast selection, we analyzed the cells segmented in one of the 7 control luciferase siRNA-treated wells of the screen. We manually annotated the 1219 objects that had been segmented by CellProfiler in this well and filtered for size (more than 4000 pixels) and nuclei content (at least 3). About one third of the 1219 objects were indeed properly segmented osteoclasts, but one third were osteoclasts surrounded by a few mononucleated cells and one third were mononucleated cell aggregates not containing osteoclasts, as exemplified in Figure 5B-D. We used this training data set to optimize osteoclast detection, using the RandomForest routine under (Breiman, 2001). The aim was to define a combination of optimal range for the geometrical and fluorescence-intensity parameters of the segmented objects calculated by CellProfiler, so as to increase the rate of true osteoclast identification. For this, we set up cross validation with 3 different random subsamples taken in the 1219-object training data set. Increasing the number of trees generated by RadomFrorest in the 3 subsamples led to convergence of the average error rate of object classification to a fixed value of 28 percent (Figure 5E). The mean accuracy of the model turned out to be 0.7616438 the 3 subsamples used, which is a good estimate of the out of sample error. The MeanRadius, FormFactor and Compactness parameters were of most importance for the prediction of object classes (Figure 5F). Applying those criterions to the 1219 objects of the training data set consistently led to an actual accuracy of 0.7361611 for the automatic selection of true osteoclasts.

To verify our model, we finally applied it to two more luciferase siRNA control wells of the screen. In each well, more than 400 osteoclasts were automatically selected among the segmented objects. Among these, we manually assessed that there were more than 80% true osteoclasts, less than 5% mononucleated cell aggregates and around 15% osteoclasts surrounded by some mononucleated cell, which is consistent with the results of the Luciferase siRNA training data set.

### Automated quantification of peripheral actin perturbation reflects visual assessment

We then applied these pipelines to the 100-test siRNA SmartPools and to the 7 control Luciferase siRNA wells of the screen for automatic selection of true osteoclasts and we measured peripheral actin intensity in the 8-pixel band. First, we calculated the median (M) and median absolute deviation (MAD) of the mean peripheral actin intensity in the 7 control Luciferase siRNA wells. Then, we measured the effect of each test siRNA (T) by calculating the Z-score as compared to Luciferase siRNA controls: (T-M)/MAD (Table 2 and Supplementary Figure 4). Comparing the results to our initial observations, we found that of the 69 siRNAs we had selected by manually as perturbing the podosome belt in that screen (Weak / Thin / Fragmented actin in Table 2), 59 did provoke a significant reduction in peripheral actin intensity as compared to control Luciferase siRNA, as shown by the negative and significant Z-scores in Table 2. This was the case for 22 out of the 28 siRNAs we had classified manually as perturbing the podosome belt in both screens (Table 1 and bold gene names in Table 2). These included Fkbp15, Spire1 and Tacc2 that we had confirmed that the siRNAs affected podosome belt organization (Figure 2) and Anillin, for which we had shown it was necessary for correct osteoclast podosome patterning and activity (Figures 3-4). Conversely, of the 31 siRNAs we visually had qualified not to affect the podosome belt (Normal in Table 2), 19 did not show a significant reduction in mean peripheral actin intensity, as shown by the positive or non-significant Z-scores in Table 2.

This suggests that our pipeline is an efficient tool for automated segmentation of osteoclasts and measurement of peripheral actin intensity, which could be useful to screen for agents capable to perturb osteoclast podosome belt and select potential inhibitors of bone resorption.

## Discussion

Osteoclasts are the major target for anti-resorptive therapies to manage osteolytic diseases. Although efficient treatments are available they suffer limitations and few patients receive appropriate treatment for osteoporosis, even after a fracture (Roux and Briot, 2018). Active research is ongoing to find novel therapeutic strategies to control pathological bone loss. In particular, we showed that targeting the podosome belt is a relevant and original strategy to prevent bone loss (Vives et al., 2015). In this context, the identification of the molecular mechanisms controlling osteoclast cytoskeleton organization is of particular interest. We provide here new candidate genes controlling osteoclast cytoskeleton. In particular, we uncover a novel non-mitotic role for Anillin. On the other hand, we established a pipeline using open source CellProfiler and R software to automatically detect actin cytoskeleton defects in the osteoclast, which could be useful for the screening of siRNAs and chemical compounds for new inhibitors of bone resorption.

Anillin is a well-documented regulator of cytokinesis, serving as a signaling platform to properly localize myosin at the cleavage furrow and coordinate RhoA activity on actin, acto-myosin contractility and central spindle microtubules (Piekny and Glotzer, 2008; van Oostende Triplet et al., 2014). Anillin was also shown to regulate neuronal migration and neurite outgrowth in *C. elegans* (Tian et al., 2015) and cell-cell junction in drosophila and human epithelial cells (Reyes et al., 2014; Wang et al., 2015). Here, we report a novel non-mitotic role of Anillin for correct podosome organization in osteoclasts. The silencing of Anillin led to improperly compacted podosomes at the cell periphery, which resulted in defective activity of the osteoclast. We did not find any changes in the level of Myosin light chain phosphorylation, tubulin acetylation nor in the localization of RhoA, phosphorylated MLC or Myosin IIA. Interestingly, we found that Anillin controls the levels of RhoA protein, which were shown to require tight regulation in order for the osteoclast to form the podosome belt and to degrade the bone (Dan Georgess et al., 2014; Ory et al., 2008). Conversely, we did not observe any effect of Anillin depletion on the levels of RhoA mRNA. Thus, Anillin controls the amount of RhoA protein in the osteoclasts, possibly through its ability to bind the GTPase as shown previously in HeLa cells (Piekny and Glotzer, 2008). The precise mechanism through which RhoA controls actin organization in osteoclasts remains poorly understood. It was reported that C3-toxin, which inhibits RhoA, RhoB and RhoC, destabilizes the sealing zone of mouse osteoclast plated on dentine (Zhang et al., 1995) or on ACC (Saltel et al., 2004). On the other hand, C3-toxin treatment of mouse osteoclasts sitting on plastic/glass favors the formation of the podosome belt (Destaing et al., 2005) but it provokes podosome disassembly in avian osteoclasts (Dan Georgess et al., 2014; Ory et al., 2008);. C3-toxin increases tubulin acetylation, via mDia2, an effector of RhoA that can activate microtubule deacetylase HDAC6. Distinct of C3-toxin treatment though, we found that Anillin siRNAs result in fewer osteoclasts with a podosome belt on plastic/glass and unchanged the levels of acetylated tubulin, as well as fewer osteoclasts with a sealing zone on ACC and reduced resorption activity. Therefore mechanisms by which C3 toxin and Anillin siRNA interfere with RhoA are distinct. It was shown that Myosin II activity was not involved in the regulation of intercellular junctions by Anilin in human epithelial (Wang et al., 2015) and we found in osteoclasts that Anillin siRNA does not affect Myosin II activity. Anillin silencing was also found to increased Jun kinase (JNK) activity in epithelial cells (Wang et al., 2015). JNK is essential downstream of RANK for osteoclast differentiation (Touaitahuata et al., 2014) and also for the formation of podosome rosettes in SrcY527F-transformed NIH3T3 fibroblasts (Pan et al., 2013), but whether JNK controls osteoclast actin cytoskeleton organization and resorption activity is unknown to date.

In order to find novel regulators of osteoclast bone resorption activity, we used a siRNA-based approach to identify novel genes controlling podosome patterning in osteoclasts sitting on plastic. The sealing zone, which only assembles on mineralized substrates, and the podosome belt (or actin ring or sealing zone like structure), which forms on non-mineralized substrates such as glass or plastic, are the most mature podosome patterns of osteoclasts. Both are made of podosomes that self assemble into a circular structure, but they differ in the density and the degree of inter-connectivity of the podosomes (Dan Georgess et al., 2014; Luxenburg et al., 2007). It was shown that podosome belt formation is regulated in a manner that parallels regulation of bone resorption (Fuller et al., 2010). Indeed, the osteoclast is able to degrade the matrix within the podosome belt (Badowski et al., 2008) and many examples illustrate that genes necessary in osteoclasts for podosome belt formation on glass are essential for bone resorption, such as Dock5 (Vives et al., 2011), Tensin 3 (Touaitahuata et al., 2016), Src (Destaing et al., 2008), FARP2 (Takegahara et al., 2010). Therefore, screening siRNAs that perturb the organization of the podosome belt on plastic/glass is a relevant strategy to identify regulators of bone resorption. Furthermore, screening for sealing zones, besides very costly, would be impossible by automated imaging because of the uneven surface of the mineralized substrates, which causes focus problems during image acquisition. Our screening was based on the observation of actin staining imaged at low magnification. The transfection of individual siRNAs for FKBP15, Spire1, Tacc2 and RalA confirmed that what we qualified in the screen as a defect in actin organization did reflect a defect in the organization of the podosomes when higher resolution imaging was performed. Therefore, low magnification imaging permits to analyze a whole well as a single image to detect actin cytoskeleton defects, which is helpful for manual and compulsory for automated screening. Ultimately though, only functional bone resorption assays can confirm the importance of a gene for bone resoption, as we show here for Anillin.

The reorganization of podosomes is a late event during osteoclast differentiation. During the differentiation mouse BMM-derived osteoclasts, precursor fusion occurs at day 2 on glass, the multinucleated cells mainly show podosome clusters and ring at day 3 and well-organized podosome belts are seen at day 4, when sealing zones are also seen on ACC. To identify regulators of podosome patterning in osteoclasts, we transfected the siRNAs at day 2 of differentiation to avoid interfering with the early differentiation processes. In fact, all 100 siRNA tested allowed the formation of large multinucleated cells where we could examine the organization of actin, suggesting that this approach is suitable to screen for genes important for the resorption process. Seeking for genes important during osteoclast differentiation would require transfection of the siRNAs earlier, for instance in BMMs before their exposition to the RANKL.

Osteoclasts are of major therapeutic interest. In the context of osteolytic diseases, they are the main targets to control pathological bone loss. But present treatments suffer limitations and the development of new treatments is necessary to respond to the increasing prevalence of osteoporosis (Roux and Briot, 2018). The quantification of osteoclasts and the analysis of their morphology are based on manual counting and visual observation, which is a time consuming and subjective process. Therefore, it is not possible to consider large scale screening to assess the effect of big libraries of chemical molecules or siRNAs on osteoclasts. We took advantage of the large amount of images we generated for our siRNA screen to set up a pipeline for automated detection of osteoclasts and objective quantification of various parameters. Osteoclasts are heterogeneous in size and shape, making automated detection uneasy. We provide here a pipeline to automatically segment the osteoclasts and calculate their geometrical parameters with the open-source resources in CellProfiler and R. We used the microtubule network for osteoclast segmentation because microtubules present the advantage over actin to distribute more evenly throughout the cell. Furthermore, whereas filamentous actin staining with phalloidin is lost when actin cytoskeleton disorganizes, on the contrary upon microtubule disorganization, such as with Siglec15 siRNAs (Figure 1B), tubulin staining resulted diffuse in the cytoplasm and thus this did not affect osteoclast segmentation. The quantification of peripheral actin intensity further proved a relevant strategy to identify siRNAs that perturb the osteoclast podosome belt; it is more reliable, experimenter independent, less time consuming, exhaustive on all segmented osteoclasts and quantitative, as compared to visual assessment.

With our pipelines, it is now possible to quantify objectively and accurately in a short time the effect of a treatment on osteoclast parameters. This procedure could also be applied for instance when screening for compounds or siRNAs that affect osteoclast differentiation. We also show that the quantification of actin enrichment at cell periphery reflects the organization of the podosome belt and thereby the capacity of the osteoclast to resorb the bone. This allows testing the effect of a treatment on osteoclast cytoskeleton to select potential inhibitors of bone resorption. With these open source pipelines, we thus provide an automated and quantitative screening strategy for treatments targeting osteoclast differentiation or function, to identify novel compounds or genes of interest in the context of osteolytic diseases.

## Acknowledgements

We acknowledge the imaging facility MRI, member of the national infrastructure France-BioImaging supported by the French National Research Agency (ANR-10-INBS-04, “Investments for the future”). We are grateful to David Guérit, Pauline Marie, Peggy Raynaud, Nathalie Morin, Cécile Gauthier-Rouvière, Franck Comunale and Mallory Genest for technical advice and reagent sharing.

## Funding

This work was funded by the French Centre National de la Recherche Scientifique (CNRS), Montpellier University and French grants from the Fondation pour la Recherche Médicale (Grant # LEQ20151134530), from the Société Française de Rhumatologie (SFR A.P. 2014 Grant # 2676) and from the GEFLUC Languedoc Roussillon (GEFLUC LR A.P. 2015 Grant # 136873-6715711) to A.B. and a grant from the Fondation ARC 2015 (Grant # PJA 20151203109) to V.V.

